# Dissecting the genetic basis of the heterosis of Y900, an elite super-hybrid rice

**DOI:** 10.1101/2022.07.16.500322

**Authors:** Zhizhong Sun, Jianxiang Peng, Qiming Lv, Jia Ding, Siyang Chen, Meijuan Duan, Qiang He, Jun Wu, Yan Tian, Dong Yu, Yanning Tan, Xiabing Sheng, Jin Chen, Xuewu Sun, Ling Liu, Rui Peng, Hai Liu, Tianshun Zhou, Na Xu, Longping Yuan, Bingbing Wang, Dingyang Yuan

## Abstract

Y900 is one of the top hybrid rice varieties with a yield exceeding 15 t/hm^2^. To dissect the mechanism of heterosis, the male parent line R900 and female parent line Y58S were sequenced using long-read and Hi-C technology. High-quality reference genomes of sizes of 396.41 Mb and 398.24 Mb were obtained for R900 and Y58S, respectively. Genome-wide variations between the parents were systematically identified, including 1,367,758 SNPs and 299,149 Indels. No megabase level structural variations exist. >75% of genes exhibited variation between the two parents. Compared with other two-line hybrids sharing the same female parent, the Geng/*japonica*-type genetic components from different male parents showed an increasing trend from phase 2-4 super-hybrid rice; Transcriptome analysis revealed that additive and dominance effects are the main genetic effects that constitute the heterosis of Y900. Allele-specific expression patterns and expression regulation patterns are quite dynamic in different tissues. For young panicle tissues, *cis*-regulation is dominant, while *trans*-regulation is more popular in leaf issues. Overdominance is more likely regulated by the *trans*-regulation mechanism. The differential gene expression and regulation pattern are closely related to Geng/*japonica* introgression. Additionally, R900 contained several excellent *japonica* haplotypes, such as *NAL1, OsSPL13, Ghd8, OsBRI1*, and *DTH2*, which make a good complement to Y58S. The fine tune mechanism through dynamic expression or regulation pattern change, especially on some key functional genes, is the base for heterosis.

## Introduction

Since the large-scale promotion and application of three-line hybrid rice in 1976, the planting area of hybrid rice has accounted for approximately 50% of the total rice area each year, and the promotion of two-line super-hybrid rice has further increased the rice yield to a new level (Cheng et al., 2007; Wu et al., 2016). The Y Liangyou 1 (Y1), Y Liangyou 2 (Y2), and Y Liangyou 900 (Y900) are representative phase 2, 3, and 4 cultivars of high-yielding super hybrid rice, respectively, and their yield potential reaches 12 t/hm^2^, 13.5 t/hm^2^, and 15 t/hm^2^. In particular, Y900 has all the characteristic traits of super high-yielding rice cultivars and is considered the culmination of 40 years of careful crossbreeding (Cheung, 2014). Although the application of hybrid rice has achieved great success, there is yet no widely accepted explanation for the molecular mechanism of their heterosis utilization.

The mechanism of heterosis has been studied in two main directions, differences in gene sequence and expression (Liu et al., 2022). High-quality genome sequences can help identify differences at the genomic level between hybrid parents. In recent years, with advancements in long-read sequencing technology, the genomes of many elite rice lines have been assembled, and the characteristics of their genome variations, including single-nucleotide polymorphisms (SNPs), insertions/deletions (Indels), structural variation (SVs), and copy number variation (CNVs), have been identified. Based on population genomic variation, a series of important heterosis-related sites, such as *Hd3a, TAC1*, and *DTH8*, have been identified through quantitative trait locus mapping or genome-wide association studies (Chen et al., 2020; Huang et al., 2016; Huang et al., 2015; Li et al., 2016; Lin et al., 2020; Zhou et al., 2012). Huang et al. constructed a quantitative trait nucleotide (QTN) map of rice and summarized the most comprehensive, key functional variations of quantitative trait genes and their roles in heterosis (Wei et al., 2021). These studies have explained the selection, fixation, and integration of dominant alleles in the process of rice hybrid breeding and provided an important genetic basis for the analysis of the molecular mechanism of heterosis. However, there are few studies on the effects of these genomic variations, especially QTN sites, on traits in the homozygous and heterozygous states, and the effects of different combinations of QTN sites that affect the same trait.

In general, compared with parental gene expression, hybrid gene expression can be divided into additive and non-additive types. Most previous studies have focussed on the importance of non-additive effects in heterosis. From the point of view of gene function, hybrids can achieve a stronger growth advantage than the two parents by integrating the advantages of the two parents in two biological pathways, cell division and photosynthesis, which are critical to plant growth and development (Liu et al., 2021). Gene expression is influenced by genomic *cis*-acting elements and *trans*-acting factors. Different epigenomic factors also affect gene expression (Ma et al., 2021). The combination of the genomic and epigenetic regulation of the hybrids as compared to the parents may lead to the transcription- and protein-level differences in different tissues between hybrids and parents. Allele-specific expression (ASE) in hybrid rice assumes that hybrids can selectively express beneficial alleles under specific conditions (Fu et al., 2022; Shao et al., 2019). Heterosis runs through almost the entire life cycle of rice, and its performance is usually tissue-dependent or temporally dynamic (Liu et al., 2022). Therefore, the exploration of the molecular mechanism of heterosis should consider the spatiotemporal specificity.

There have been many studies on the mechanism of heterosis in hybrid rice in the areas of genome assembly, variation mining, heterotic site identification, and gene expression and regulation. However, the molecular mechanism of heterosis formation has not been systematically and comprehensively investigated in the two dimensions of genomic variation and gene expression. Based on assembling the genomes of Y58S and R900, variations at the whole-genome level were identified and beneficial Geng/*japonica* (GJ) haplotypes carrying several important functional genes were found in the male parent R900. The differentially expressed genes (DEGs) in four tissues between hybrid and parents were checked, and the relationship between *cis-*/*trans-*regulation and heterosis in rice was investigated for the first time. Finally, the effects of several important loci in the F2 population were verified. This study provides new insights for the systematic interpretation of the basis of heterosis formation in rice.

## Results

### Phenotypic analysis and two high-quality reference genome assemblies

Y900 outperformed its parents in total grain number per plant (GNPP), panicle length (PL), grain length (GL), and plant height (PH), especially in the grain yield per plant (GYPP) (43.9 g) exceeding the male parent (28.7 g) by approximately 52.96%, showing superior yield heterosis (Figure 1a and Figure 1b; Table S1). Y1 and Y2 showed consistent heterosis with Y900 in most traits, i.e., heterosis over both parents in GNPP, PL, and PH and heterosis over one parent in the effective panicle number (EPN), grain number per panicle (GNPPa), thousand-grain weight (TGW), GYPP, heading date (HD), and flag leaf width (FLW). Consistent with the hybrid rice LYP9 (Li et al., 2016), all three hybrids took full advantage of over-male-parent heterosis (OMPH) in EPN and over-female-parent heterosis (OFPH) in GNPPa (Figure 1c; Table S1), which may be important for the heterosis in the yield of the two-line hybrid rice. Although Y900 was inferior to the male parent in GNPPa, it exhibited the highest utilization rate of OMPH in EPN (73.91%) and OFPH in GNPP (16.17%), which made the GYPP of Y900 exhibit heterosis over both parents, with an increase of 31.44% and 12.85% over that of Y1 (33.4 g) and Y2 (38.9 g), respectively (Figure 1c).

**Figure 1.**
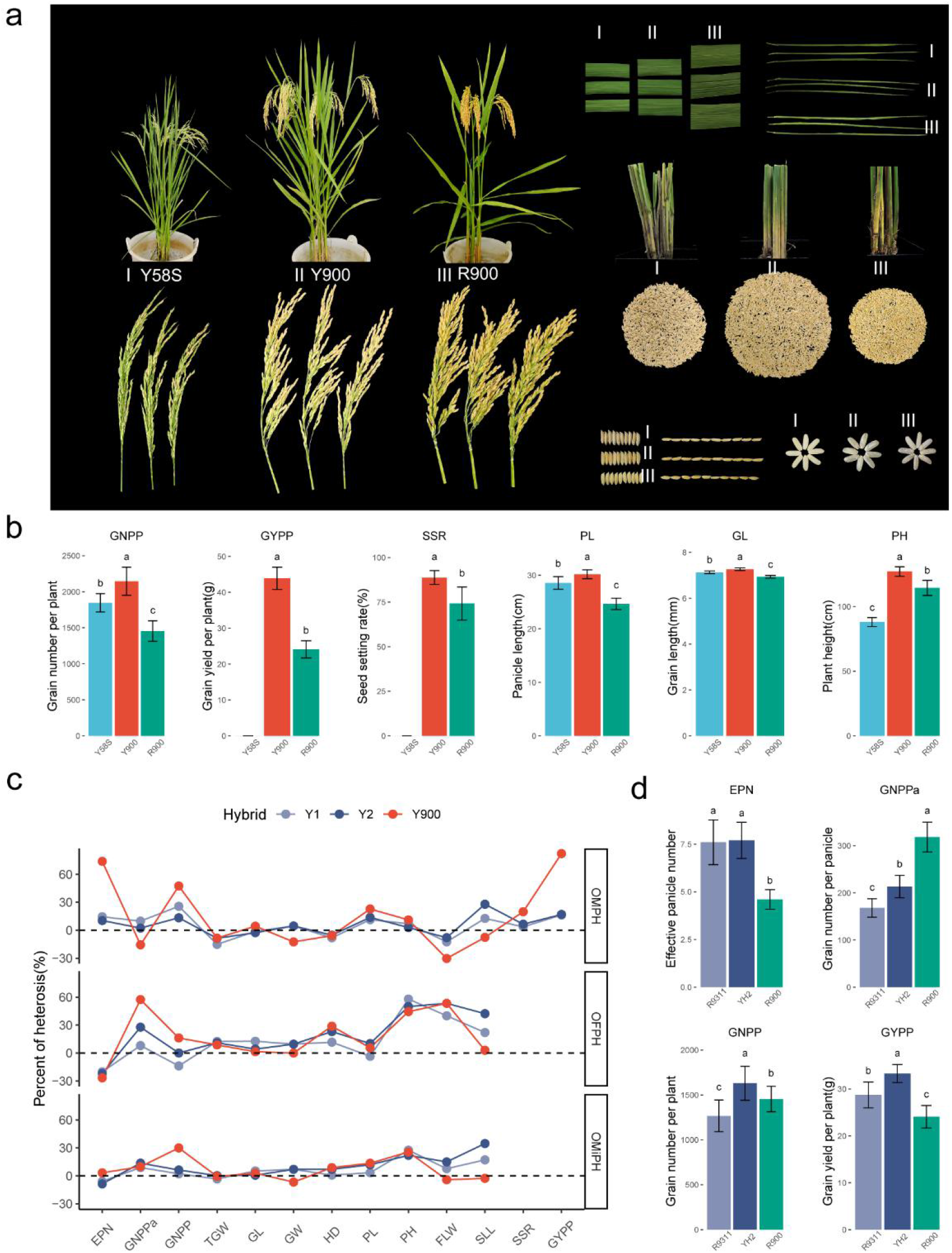
Phenotypic heterosis in super-hybrid rice Y1, Y2, and Y900. **a** Comparison of plant, panicle, leaf, and grain morphology of Y900 and its two parents, R900 and Y58S; **b** Heterosis of Y900 over both parents in yield-related traits; **c** Comparison of heterosis rates for 13 important agronomic traits in Y1, Y2, and Y900. OMPH, OFPH, and OMiPH represent the over-male-parent heterosis, over-female-parent heterosis, and over-mid-parent heterosis, respectively; **d** Comparison of yield factors for R9311, YH2, and R900. Different letters above the bars in b and d represent significant differences by one-way ANOVA (*P* < 0.05).

Compared with the male parents R9311 and Yuanhui 2 (YH2), respectively corresponding to Y1 and Y2, the restore line R900 had the most compact plants with shorter PH and larger FLW (Figure 1a; Table S1). Although R900 was slightly inferior in EPN, GNPP, and GYPP, the GNPPa of R900 was much higher than those of R9311 and YH2. In addition, the HD of R900 is the longest, nearly 12 days more than that of R9311 (Table S1). These characteristics provide more time for the growth and development of hybrid rice Y900 and lay the foundation for lodging resistance and high fertilizer tolerance, which are conducive to superhigh yield. Whereas R900 is perfectly complementary with the yield factors of Y58S (smaller GNPPa, earlier HD, and more EPN), ultimately resulting in strong heterosis with a higher yield for hybrid rice Y900 than Y1 and Y2 (Figure 1a; Figure 1c; Table S1).

The genomes of R900 and Y58S were assembled using ~110 Gb and ~101 Gb long reads generated by PacBio Sequel II (Figure S1), and after polishing with ~85 Gb and ~75 Gb short reads (Table S2), we obtained assembled sequences of ~396 Mb and ~398 Mb in size, with contig N50s of ~16 Mb and ~17 Mb, respectively. Ultimately, ~98% (389.04 Mb and 389.54 Mb) of the sequence were anchored onto 12 chromosomes using ~45 Gb and ~46 Gb of Hi-C sequencing data, with gaps of 42 and 35, respectively (Figure 2a; Figure S2; Table 1; Table S4). Over 99% of the next-generation sequencing (NGS) reads could be properly aligned to both genomes, and the benchmarking universal single-copy orthologues (BUSCO) were 98.70% and 98.60%, the highest among the known important rice reference genomes (Figure S3). In addition, the high long terminal repeated assembly index (LAI) (>23) and the very strong colinearity with genomes such as NIP, R498, and R9311 indicate that the quality of both genomes is excellent (Table 1; Figure S4; Table S6). After masking ~54% of the repetitive sequences and removing transposable element (TE)-related genes, we predicted 40417 and 41583 protein-coding genes and 667-1988 non-coding RNA genes in the R900 and Y58S genomes, respectively. A total of 89.81% and 87.42% of the protein sequences were annotated in the functional database (Table S7-S10).

**Figure 2.**
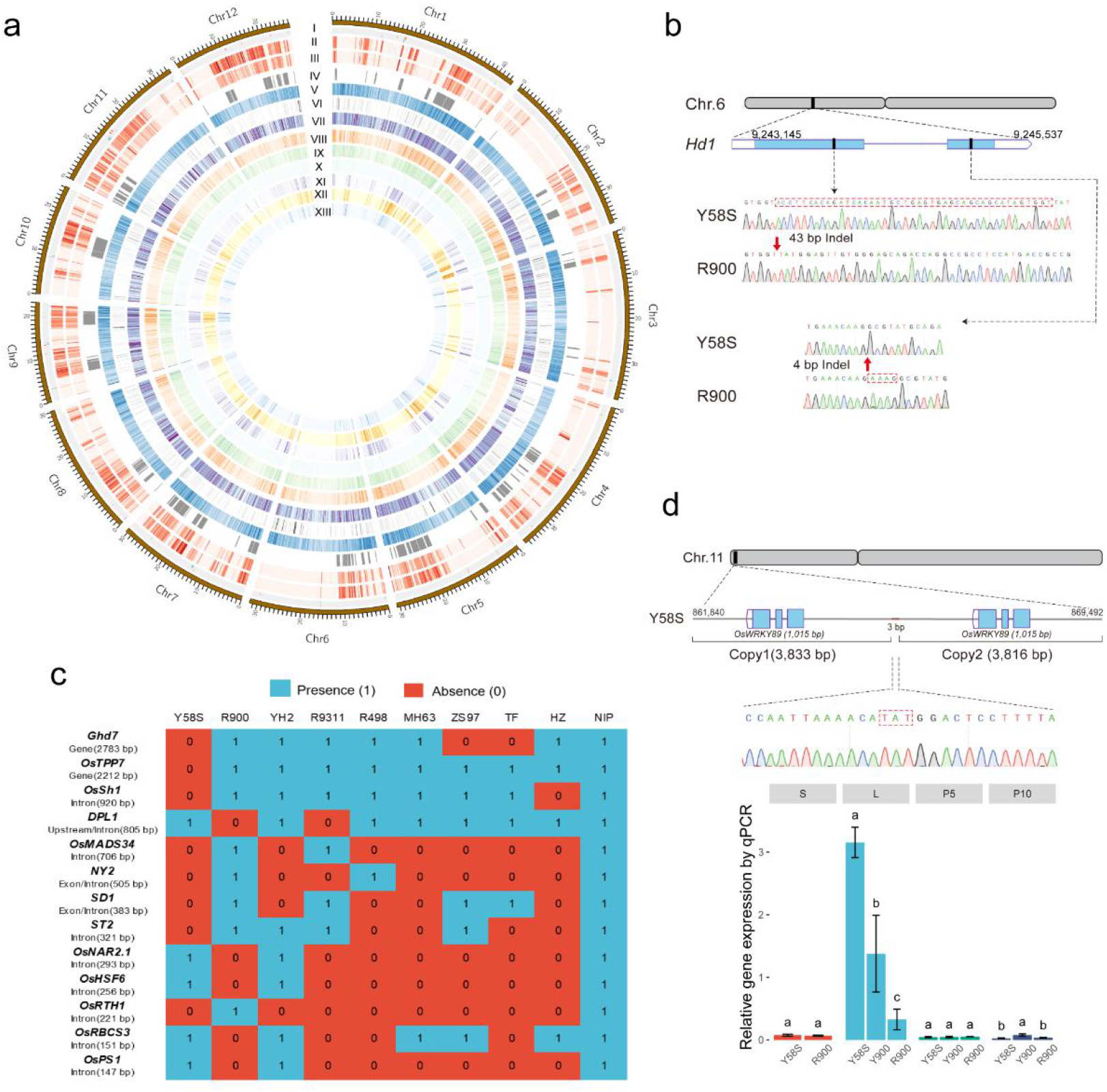
Y900 biparental genomic features and variations. **a** Genome circos map. I: Distribution of gaps in Y58S (red) and R900 (blue); SNP and Indel densities in Y58S(II) and R900(III); IV: Regions of variations between the Xian/*Indica* and Geng*/Japonica* parents; V: Gene density in Y58S; VI: Density distribution ratio of the cloned genes of parental variations (2,790) to all variant genes (36,477); VII: Density distribution ratio of parental DEGs (13,490) to all expressed genes (36,393); VIII-X: Densities of genes with effects of PDO, DO, and ODO; XI-XIII: Densities of genes with *cis, trans*, and both *cis* and *trans* expression regulation patterns. Colour shades represent the magnitude of the distribution density; **b** Identification and Sanger sequencing verification of two Indels of *Hd1*; **c** PAVs of the cloned genes in important rice reference genomes; **d** CNV identification, Sanger sequencing, and expression by qPCR in different tissues of *OsWRKY89*; different letters above the bars represent significant differences by one-way ANOVA (*P* < 0.05).

**Table 1.** Summary of the assembly and annotation features of R900 and Y58S.

|  | R900 | Y58S |
| --- | --- | --- |
| Genome coverage (x) | 273.98 | 251.95 |
| Assembly length (Mb) | 396.41 | 398.24 |
| Chromosome anchoring rate (%) | 98.14 | 97.82 |
| Number of contigs | 230 | 177 |
| Contig N50 (Mb) | 15.95 | 16.59 |
| Largest contig (Mb) | 34.76 | 33.28 |
| Scaffold N50 (Mb) | 31.86 | 31.81 |
| Number of gaps | 42 | 35 |
| Genome BUSCO (%) | 98.70 | 98.60 |
| LAI score | 23.73 | 23.90 |
| Mapping rate (%) | 99.83 | 99.89 |
| GC content (%) | 43.71 | 43.71 |
| Repeat content (%) | 54.27 | 54.82 |
| Number of non-TE genes | 40, 417 | 41, 583 |
| Protein BUSCO (%) | 95.40 | 95.60 |

### Genome-wide variation and GJ introgression differentiation

A total of 1,367,758 SNPs and 299,149 Indels (149,604 insertions and 149,545 deletions) were identified between R900 and Y58S genomes, which were distributed in similar densities (Figure 2a). The number of variants between R900 and Y58S was almost minimal compared to the assembled rice genome (Figure S5 and S6; Table S11). At least 36,477 genes (76.49%) had sequence differences between R900 and Y58S, including 2,790 cloned genes (75.32%) (Data S1). The distribution pattern of these varied cloned genes was not always consistent with the genome-wide distribution pattern of SNP/Indel. An example can be found on the long arm of Chr6, where many cloned genes showed variation between the two parents, but the SNP/Indel density was relatively low (Figure 2a). More importantly, among 342 QTN sites of 253 genes (Wei et al., 2021), 59 QTN (17.25%) of 47 genes (18.58%) differed between the two parents of Y900, e.g. *Hd1* and *NAL1* (Data S2; Figure S7; Figure S8). The QTN for *Hd1*, located in the Y58S genome at Chr6: 9,245,074, is C in Y58S and CAAGA in R900 (as in GJ) and delays flowering. Moreover, a 43 bp deletion of *Hd1* in R900 was found, which resulted in a shift mutation, and the effect was first identified similarly to the 4 bp QTN above (Figure 2b; Data S12).

Similarly, the differences in PAVs, translocations and inversions between R900 and Y58S were not significant among the comparisons (Figure S9; Table S11). Although no megabase level variations were found, some important functional genes were identified in PAVs, and more PAVs were located upstream of 2kb, introns and exons of the gene (Figure 2c; Figure S10; Figure S11; Figure S12). *Ghd7* is completely absent in the sterile lines Y58S, ZS97 and TF, while it is present in restorers such as R900, indicating that the complete deletion haplotype of *Ghd7* plays an important role in the two-line hybrid rice selection system. Simultaneously, 687 CNVs with 321 copy gains and 366 copy losses were identified respectively (Table S11; Data S3). One of the CNVs on the short arm of chromosome 11 (Y58S Chr11: 861,840-869,492) was first identified, and *OsWRKY89* (OsY58Sg08701) within this fragment is a transcription factor that affects early plant growth and development (Wang et al., 2007). Further analysis revealed two copies of this gene in Y58S and Lemont, but only one copy in R900 and the sterile line PA64S. Lemont and PA64S are parents in the Y58S breeding lineage, so the copies probably originated from Lemont. Gene expression analysis showed that *OsWRKY89* was much more abundant in flag leaf (L), and the expression of Y58S and Y900 was significantly higher than that of R900, showing a dominant effect (Figure 2d).

After constructing XI-GJ genetic compositions and introgression bin maps, we found some hotspot regions of GJ introgression were relatively consistent with the hotspot regions of variation between their genomes, such as Chr5: 17.9-27.3 Mb, Chr7: 22.3-27.1 Mb in Y58S, and Chr1: 29.2-34.4 Mb, Chr9: 17.7-21.2 Mb in R900 (Figure 2a; Figure S5; Figure 3a). The XI component (XI-ind + XI-aus + XI-admix) of Y58S, R900, YH2 and R9311 were all above 60% and the GJ components (GJ-admix + GJ-tmp + GJ-trp) were ~18.37%, ~14.31%, ~3.40% and ~2.11%, respectively (Figure 3b). Notably, the GJ introgression fragments were most abundant in Y58S (~68.7 Mb), presumably due to the presence of Lemont and Paddy in its breeding lineage. R9311, YH2, and R900 had sequentially higher GJ introgression fragment sizes (~7.9 Mb, ~12.7 Mb, and ~53.5 Mb), indicating a degree increase in the use of hybrid heterosis between XI and GJ in phase 2-4 super rice (Figure 3b). Of the 3,704 rice cloned genes, 3,469 (~94%) could be assigned to the bin maps, Y58S (729) and R900 (626) had far more cloned genes in the GJ region than YH2 (103) and R9311 (118) (Figure 3a). Among the combinations of genetic components, Y58S as GJ-admix and the other three varieties as XI-ind combinations had the most cloned genes (127). This was followed by R900 for GJ-admix (112) and GJ-tmp (56), indicating that R900 utilized more known GJ loci relative to YH2 and R9311 (Figure 3c). Except for *ALK* and *DPL2*, which selected XI genotype, 51 QTN sites for the other 36 genes of Y58S were located in the GJ region (Figure 3a). R900 (61 QTN sites for 40 genes, with only 4 loci inconsistent with NIP) was much higher than YH2 (22 QTN sites for 13 genes) and R9311 (17 QTN sites for 8 genes) in the proportion of GJ alleles exploited (Table S15). In particular, the QTN of 11 genes, including *OsSPL13, NOG1, D61, bZIP73, TAC1*, and *DTH2*, were only GJ in R900 (Figure 3a), which may be an important basis for the strong heterosis of Y900.

**Figure 3.**
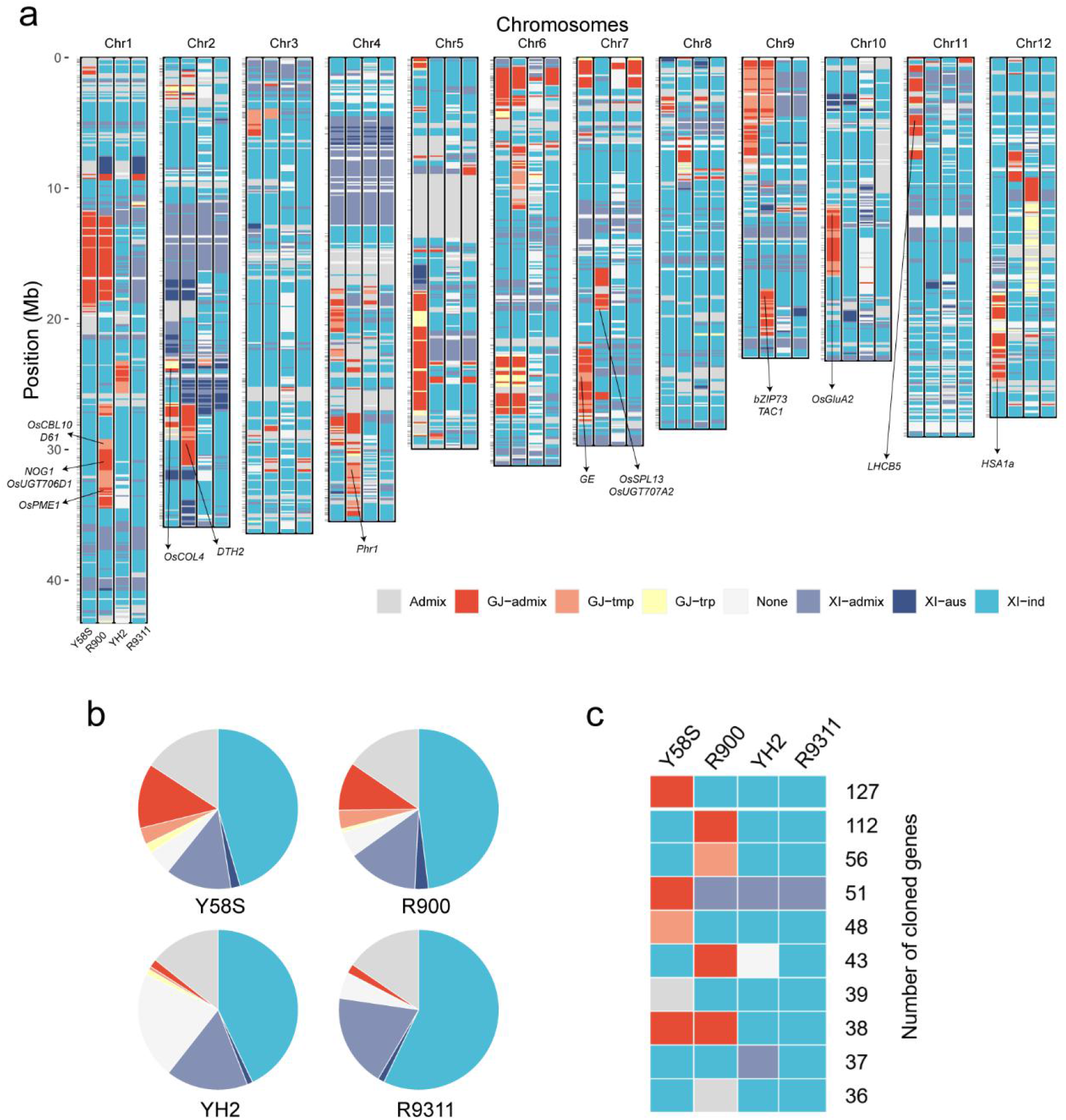
Genetic composition and introgression of super-hybrid rice Y1, Y2, and Y900 parents. **a** Genetic composition distribution of Y58S, R900, YH2, and R9311. The short horizontal line to the left of each chromosome shows the locations of 3,704 cloned rice genes; from left to right, the binmaps show Y58S, R900, YH2, and R9311; **b** Proportions of genetic components from the parents; **c** Ranking of the number of parental genetic component combinations in cloned genes (top 10).

### Transcriptome analysis of Y900 and its parents

The transcriptomes of four tissues of Y900 and its parents, including stem (S), flag leaf (L), 5 mm (P5) and 10 mm length young panicle (P10) were analyzed and the data showed good reproducibility (Figure S13). A total of 36,393 genes were expressed in all tissues (Table S16), of which 25% (9,061) genes were of low expression (TPM <= 1). Approximately 70% (25,689) of the genes were expressed in all four tissues, and 30% of the genes were tissue specific. We identified 2,139 variety-specific genes and found that the number of hybrid-specific genes expressed in L was the highest (196) (Figure S14; Table S17), most likely due to transcriptional regulators of one parent acting on the promoter of the other parent. This crossed trans-regulatory mechanism, although to be further validated, should be a component of heterosis. Analysis of differentially expressed genes (DEGs) revealed that about 37% (13,490) of the genes differed among the three varieties. The number of DEGs in the young panicle was significantly lower than in the stem and leaf, and the number of DEGs between parents in the leaf was as high as 7,180 (53%) (Figure 4a). It is easy to understand that the gene expression differences between varieties are relatively small in the early stage of tissue development, while during and after tissue establishment, the gene expression differences between varieties increase and become fixed, resulting in phenotypic differences between varieties. In L, P5, and P10, the DEGs between hybrids and parents were less than those between parents, and the DEGs between hybrids and parents were less than the DEGs. But in S, we unexpectedly found that the DEGs between hybrids and either parent were 40% more than those between parents (2,475), and the DEGs between hybrids and parents (3,951) were more than the DEGs between hybrids and parents (3,490), suggesting that over-parent heterosis is more likely to occur in S of the hybrid.

**Figure 4.**
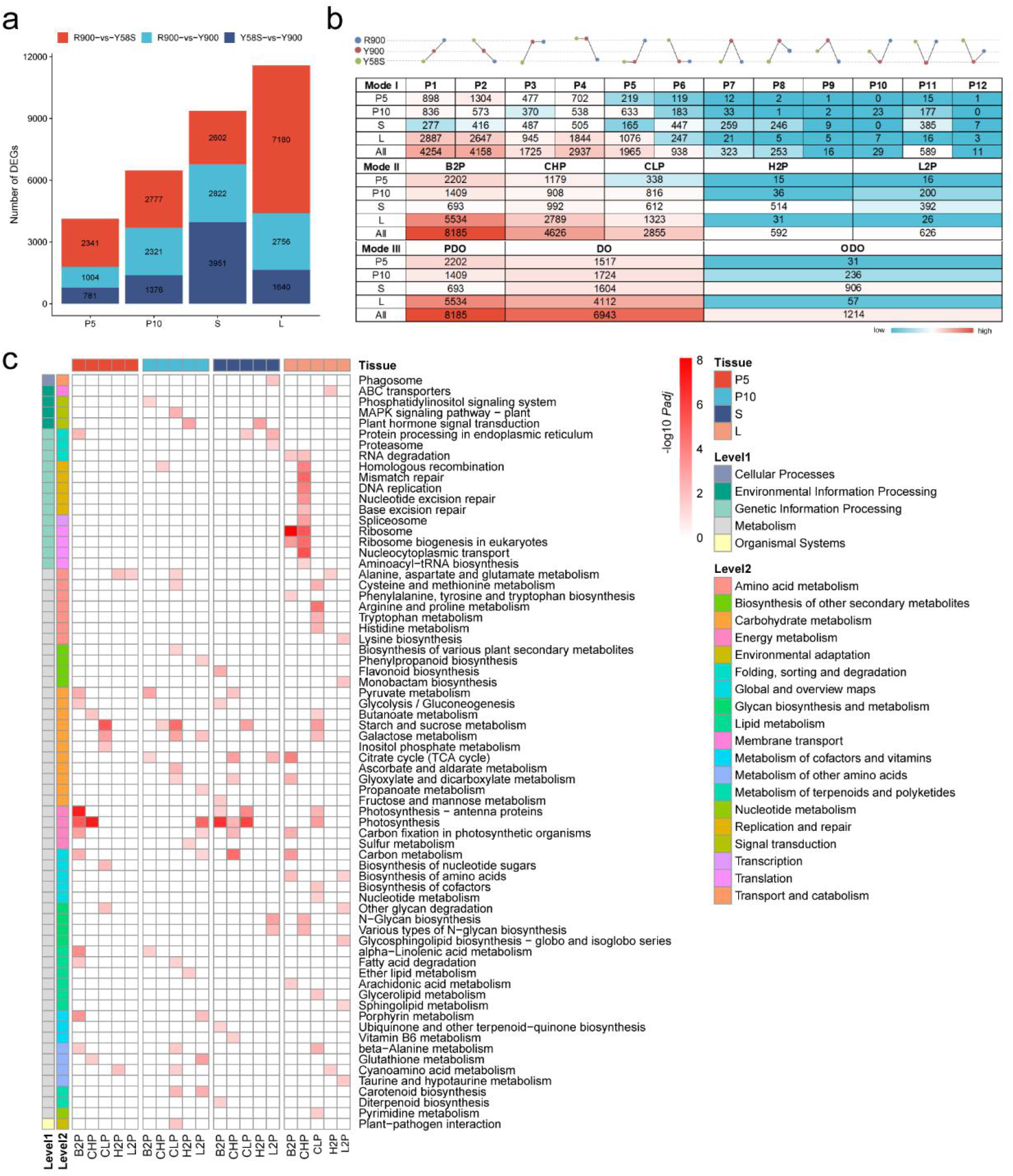
Multi tissue transcriptome analysis of Y900 and its parents. **a** DEGs in four tissues of R900, Y58S, and Y900; **b** Division of the gene expression modes. There are 12, 5, and 3 modes in Mode I, Mode II, and Mode III, respectively. The shades of red and cyan represent the numbers; **c** KEGG pathway enrichment analysis of Mode II genes. The line labels Level1 and Level2 represent two KEGG levels, and color shades indicate the degree of enrichment (*P adjust* < 0.05).

The proportion of DEGs shared in common by the three groups was only 5.8-12.7% (Figure S15), indicating that the gene expression in different tissues of the three varieties is quite different. By further analyzing the differences in the expression levels of these genes between the hybrids and their parents, we found that the expression levels of 1,214 genes in the hybrids were significantly higher or lower than those of the parents (ODO mode), the expression levels of 6,943 genes in the hybrids are similar to one parent but significantly higher than the other parent (dominance (DO) mode), and the expression levels of 8,185 genes in the hybrid were between the two parental levels (partial dominance (PDO) mode) (Figure 4b; Data S7). Thus, the contribution of ODO to heterosis may not be as strong as the additive and dominance effects, which differs from the previous view that the genetic pattern of hybrids is mainly ODO, probably due to the use of different methods and varieties (Chen et al., 2018; Fu et al., 2022). Distributions of the DEGs and different expression models were largely consistent with the distribution of SNP and/or Indel (Figure 2a). The gene expression patterns in different tissues were significantly different. Although there were many DEGs in the L, most of them were in the PDO and DO modes, and only 0.5% (57) were in the ODO mode. In the young panicles at the two different stages, the proportion of the ODO mode increased to 0.8% and 7%, respectively. Consistent with the above conclusion, the proportion of ODO in S was as high as 28%, which was even higher than the proportion of PDO.

Analysis of the gene regulatory network during the development of *Arabidopsis thaliana* seedlings has shown that the hybrid can achieve a growth advantage over the two parents by integrating their respective strengths in photosynthesis and cell division, two biological pathways essential for plant growth and development (Liu et al., 2021). In this study, KEGG enrichment analysis showed that many DEGs in S and young panicles were related to photosynthesis, mainly showing an additive effect or a dominance effect (over-parent and below-parent), but in P10, the expression levels in the tissues were lower than those of the parents (Figure 4c). The close-to-lower-parent (CLP) genes in all tissues were enriched in starch and sucrose metabolism. The expression levels of genes related to plant hormone signal transduction in S and P10 were over the parents. In contrast, there were relatively few DEGs related to photosynthesis in L, and many genes were related to cell division or material metabolism (Figure 4c). These results indicated that in the strong dominant hybrid combination, the photosynthesis efficiency between different parental leaves might not be significantly different (all already high), but in hybrids, the efficiency of basic cell activities (over-parent) increases, the metabolism of related substances (below-parent) decreases, and the efficiency of photosynthesis increases in other tissues (S), thus achieving overall heterosis.

### ASE in Y900

To confirm that the alleles of DO and ODO gene expression in hybrids are mainly from the male or female parent, 7,562 allele-specific genes with differences between the parental alleles were identified (SNP was present in the transcript). Analysis of the expression in the four tissues revealed a total of 3,048 (40%) allele-specific expressed genes (ASEGs), and the ratio of genes that leaned more towards the male parent to those towards the female parent was close to 1:1 (Figure 5). We identified several potentially imprinted genes with only one allele expressed in F1, where the genes in P5 are adjacent to each other and only the Y58S allele is expressed (Table S12). Therefore, in most cases, the two alleles from the parents can be expressed simultaneously in the hybrids, 40% of the genes in some tissues are biased towards one of the parents, and very few genes show complete dominant or recessive expression.

**Figure 5.**
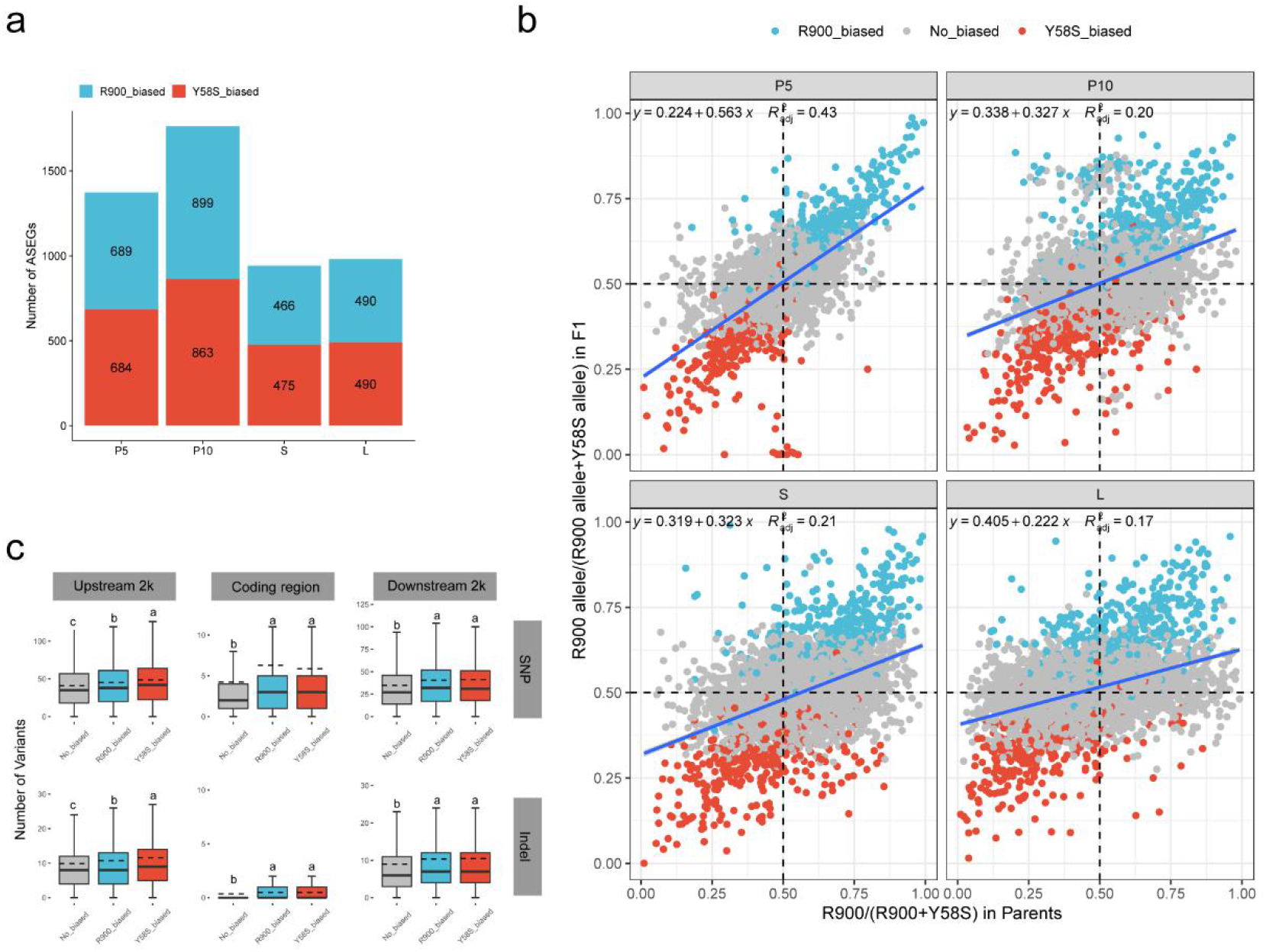
ASE in Y900. **a** Bias of ASEGs in different tissues; **b** Relationship between ASE in F1 and the expression in parents. The figures show the linear regression equations and corrected coefficients of determination R^2^ for each tissue; **c** Relationships between the numbers of SNPs and Indels and ASEG. Different letters above the bars represent significant differences by one-way ANOVA (Duncan’s multiple range test, *P* < 0.05).

Further study found that the expression of ASEGs in different tissues changed dynamically. Among the four tissues, the number of ASEGs was the highest in young panicle P10 (1,763), accounting for 27% of the ASEGs in this tissue and approximately 20% in the other three tissues. The fold difference in the allele expression in tissues of young panicles P5 and P10 was smaller than that in S and L (Figure S16), a trend that was consistent with the differential expression of related genes in both parents, indicating that both copies from both parents could be fully expressed in the tissues of actively growing hybrids. With the differentiation of tissues, the bias of ASEGs also changed. Among all the tissues, only 122 had the same parental bias, and the ratio of male-parent bias R900 (61) to female-parent bias Y58S (61) was exactly 1:1 (Figure S17). There were 2,420 genes with inconsistent bias, including 115 genes with opposite biases in two tissues and 2,305 genes with a certain bias in some tissues but no bias in other tissues (Data S8), indicating that similar to the three-line hybrid rice (Shao et al., 2019), there are far more inconsistent ASEGs than consistent ASEGs in two-line hybrid rice. Given that the *cis*-elements of genes in different tissues are the same, these dynamic changes in bias reflect the fine regulation of gene expression by transcription factors from parents in specific tissues.

This study compared the biases of ASEGs with the differences in gene expression between the parents and found that most ASEGs (74-89%) had a good correlation with the differences in gene expression between the parents (Figure 5b). The correlation was the highest in P5, while the correlation is lower in L. On the one hand, these data indicate that the differences between the parents were relatively well reflected in the hybrids, which is the basis for heterosis formation, and these genes may be subjected to relatively strong *cis*-regulation; on the other hand, these data indicate that the role of *trans*-regulation in mature tissues is greater than that in young tissues. This study further compared the variations in the vicinity of alleles between the parents and found that ASEGs had significantly more SNPs and indels than non-ASEGs regardless of whether the SNPs and Indels were in the upstream 2 kb (promoter region), the coding region, or the downstream 2 kb (Figure 5c), demonstrating the role of *cis*-regulatory elements in ASEGs. In the aquatic plant *Nelumbo nucifera*, ASEGs in the promoter region are more active than the conserved genes (Gao et al., 2021).

### Relationship between gene expression regulatory mechanism and expression pattern

To further investigate the regulation of the expression of 7,562 allele-specific genes, we compared the differences in the expression of these genes between parents (A) and the differences in the expression of alleles from the male and female parents in hybrids (B), and divided the expression regulation patterns of these genes into seven categories (I-VII) as described (Bao et al., 2019) (Figure 6a; Data S9). Overall, the expression of most alleles in young panicles did not differ between parents or between two copies in hybrids (conserved, 51.48-61.74%) or was difficult to determine (ambiguous, 22.08-30.72%), and most of the remaining alleles were *cis*-regulated (*cis* only: 12.61-14.25%). However, in S and L, the number of conserved genes was significantly reduced, the number of *cis*-regulated genes was also reduced to approximately 8-9%, and the number of *trans*-regulated genes was significantly increased. Especially in the L, the gene expression regulatory mechanism was very different. Approximately 29.74% of the genes were *trans*-regulated only, which was even higher than the proportion of the conserved genes. The expression levels of many expressed genes in L differed significantly between the parents, but the expression levels of the two copies in the hybrids were almost the same (Figure 5b; Figure S18). The number of genes regulated by categories III-V was relatively small and was mainly concentrated in S and L (Figure 6a). In this study, the proportion of *trans*-only regulation was significantly higher than that in a study of cotton (0.9-1.1%) (Bao et al., 2019), which may be related to the cotton tissues used (ovules at 10 and 20 d post-anthesis).

**Figure 6.**
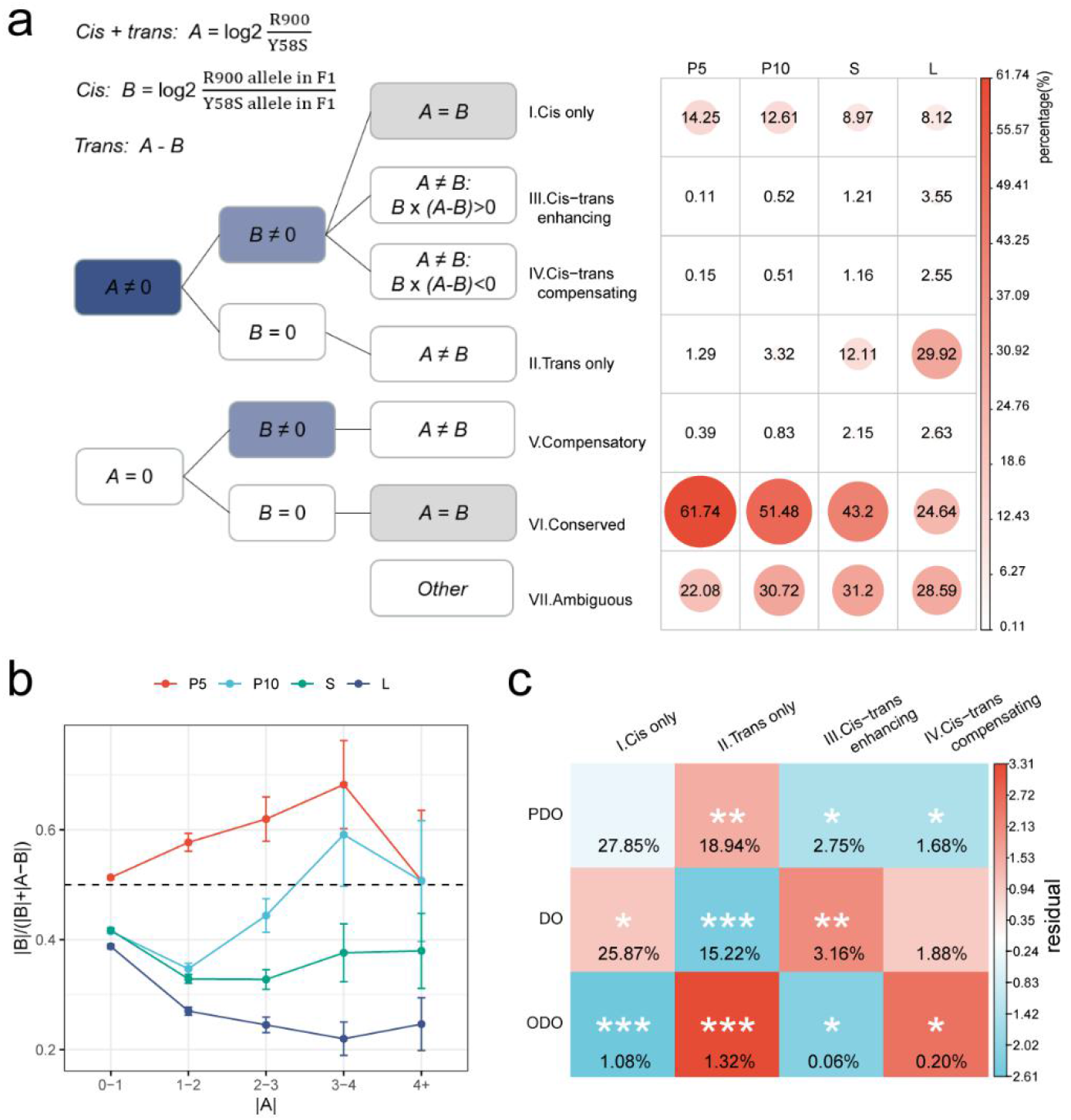
Relationship between *cis-/trans-*regulation patterns and modes of gene expression. **a** Classification and identification of expression regulation patterns of alleles in F1 and parents. Differential gene expression between the parents reflects *cis-* and *trans-*acting effects (A), while differential expression of alleles from the two parents in F1 reflects the *cis* effect (B). The *trans* effect cannot be indirectly measured but can be calculated using A-B. Gene expression regulation can be classified as categories I-VII based on the comparisons of A and 0, B and 0, and A and B. The size and color of the circles represent the proportion of the regulation category in a particular tissue; **b** Proportion of the contribution of *cis-*regulation to the expression differences in the parents. The x-axis represents absolute differences in gene expression between the parents, while the y-axis represents the proportion of differential expression caused by the *cis* effect relative to that caused by the total effect. Error lines indicate the 95% confidence intervals; **c** Relationship between modes of gene expression (rows) and *cis-/trans-*regulation patterns (columns). The color indicates the residual size calculated by an independent Pearson’s chi-square test; positive residuals represent enrichment in the gene number (red), i.e., more genes than expected under an independent null model, and negative residuals represent depletion in the gene number (cyan), i.e., fewer genes than expected under an independent null model. Statistical significance was tested by Fisher’s exact test, and *, **, and *** represent *P* values of less than 0.05, 0.01, and 0.001, respectively.

The gene regulation pattern changes dynamically in different tissues. In the four types of tissues, 3,421 genes were identified in seven categories. Except for the ambiguous category, only 201 genes had the same regulatory pattern (constitutive regulation), which were mainly the *cis* only and conserved categories (Figure S19). Most genes may adopt different regulation patterns in different tissues, and the expression differences of *trans*-regulated genes in different tissues were more obvious, which is consistent with the dynamic changes in the gene expression regulation in different tissues in maize (Zhou et al., 2019). The number of SNPs and Indels in genes regulated in *cis* only was significantly higher than that in genes regulated in *trans* only or no-divergence genes (including compensatory and conserved), and it was not significantly different between *cis* and *trans* (including enhancing and compensating) in the upstream and downstream regions (Figure S20).

When the difference in allele expression between parents was not significant (0-to 2-fold), the contribution of *cis*-regulation to gene expression in each tissue was 40-50%. When the difference was between 2-to 16-fold, the greater the difference in the allele expression in young panicles, the greater the role of *cis*-regulation, and the highest contribution was found in the P5, up to 70% (a difference of 8-to 16-fold). In L, the contribution of *cis*-regulation gradually decreases to approximately 20% at the lowest (difference of 8-to 16-fold) (Figure 6b), indicating *trans*-regulation makes a greater contribution than *cis*-regulation in mature tissues. Among the seven regulatory categories, categories I-IV had the greatest impact on the difference between parents, especially *cis*/*trans* (Figure S21). In cotton (Bao et al., 2019) and chili (Diaz-Valenzuela et al., 2020), a similar phenomenon has been found.

The relationship between gene regulation and heterosis was studied, and the results showed that when there was a significant difference in the expression of parental alleles (A ≠ 0), there was a significant correlation between the two (Figure 6c). The proportion of PDO genes regulated in *cis* only was the highest (27.85%), which was consistent with expectations. However, the proportion of DO genes regulated in *cis* only (25.87%) was slightly higher than expected, while the proportion of corresponding ODO genes was extremely significantly lower than expected. In contrast, the proportion of DO genes regulated in *trans* only (15.22%) was significantly lower than expected, but the proportion of ODO and PDO genes regulated in *trans* only was extremely significantly higher than expected (1.32%), indicating that in rice heterosis, the mechanism of *trans-*only regulation plays a key role in the ODO mode (ODO genes) and the additive effect mode (PDO genes), while the contribution of the regulatory mechanism of *cis* only to the dominance mode (DO genes) is higher than expected.

### Verification of the heterosis effect of different genotypes

Among all 3,704 cloned functional genes in rice, 1,854 were DEGs between the parents and the hybrid and could be divided into the expression patterns of Mode I-III (Data S10), and the expression patterns of 33 genes in four tissues could be identified (Figure S22). For 1,110 genes, their ASE and gene regulation (categories I-VII) could be determined (Data S11), and for 739 functional genes, both the expression pattern and expression regulation category were cleared. The relative proportions of the expression patterns to regulation categories of these functional genes in various tissues were consistent with the overall proportions of all genes, indicating that the expression regulatory mechanisms are relatively representative.

Subsequently, we checked some important loci related to heterosis, such as *Ghd8/RH8*, which is the main heterosis gene of two-line hybrid rice (Huang et al., 2015; Li et al., 2016). The Y900 (heterozygous genotype of *Ghd8*) exhibited significant heterosis over both parents on the traits of GNPP and GNPP, and GYPP also showed significant heterosis over the higher parent (13.6%) (Figure 7a). The expression of *LAX1* in P5 was higher than that in other tissues, showing a DO mode between the hybrid and the parents. According to the genotyping of QTN in the population, when the gene was in the heterozygous state, traits, such as PL, GNPP, and yield, showed significant heterosis over the male parent, and the SSR showed significant heterosis over the female parent (Figure 7c). *NAL1* was highly expressed in young panicles and was only biased in P10 of hybrids (biased toward R900) and regulated by cis-acting (Data S11). The FLW of the homozygous paternal genotype (2.48 cm) was extremely significantly greater than that of the maternal genotype (2.04 cm), and the heterozygous genotype was biased toward the paternal genotype (Figure 7d). The alleles from R900 could significantly improve FLW, GNPP and PH, and increase yield heterosis.

**Figure 7.**
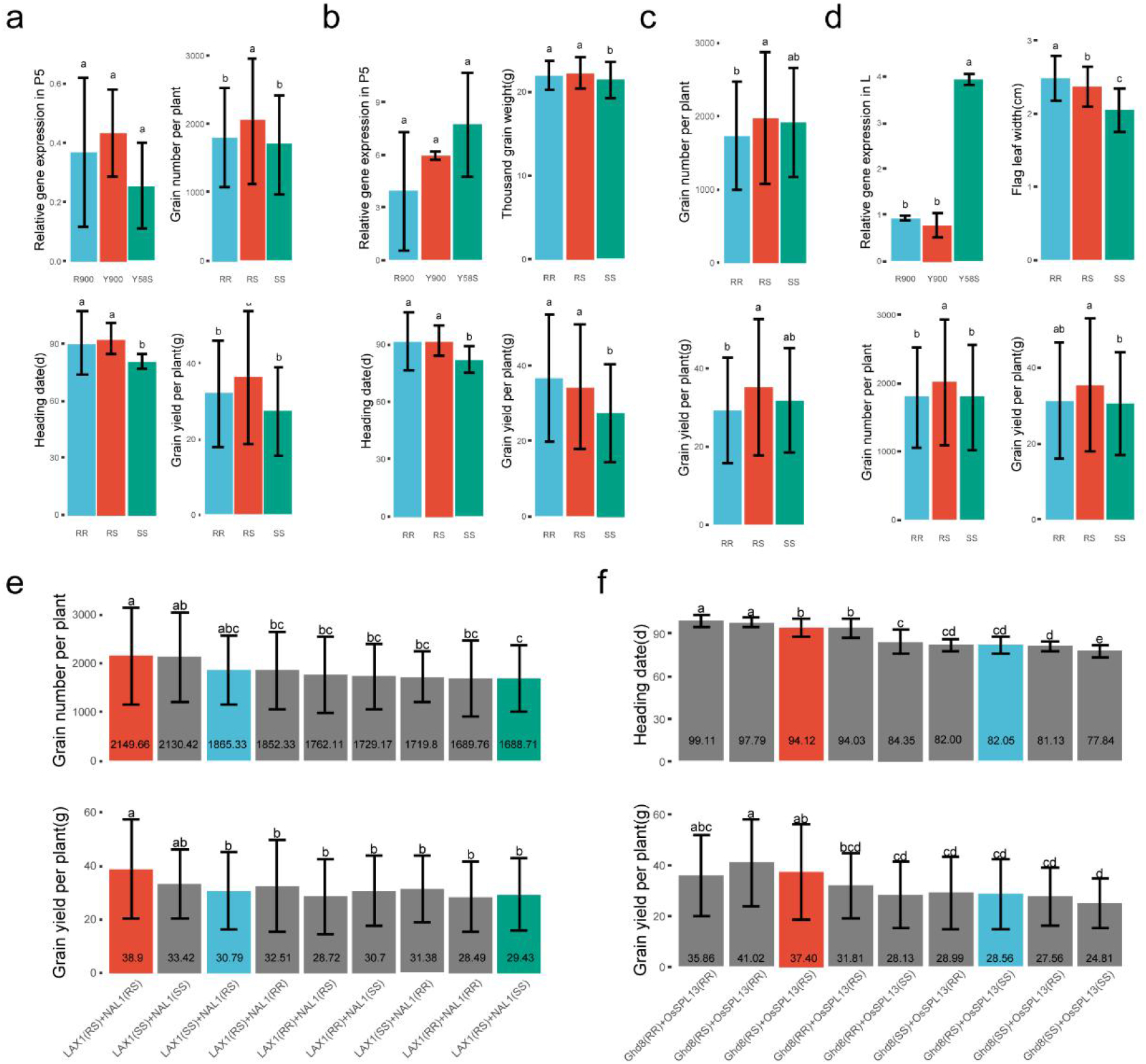
Effects of different heterosis loci. **a** Verification of *Ghd8* expression in P5 and comparison of the three different genotypes in terms of GNPP, HD, and GYPP. The three genotypes, RR, RS, and SS, correspond to homozygous R900, heterozygous Y900, and homozygous Y58S, respectively; **b** Verification of *OsSPL13* expression in tissue P5 and comparison of the three different genotypes in terms of TGW, HD, and GYPP; **c** Comparison of the three different genotypes of *LAX1* in terms of GNPP and GYPP; **d** Expression verification in flag leaf L and comparison of the three different genotypes in terms of FLW, GNPP, and GYPP; **e** Comparison of the nine different genotypes of *LAX1* and *NAL1* in terms of GNPP and GYPP. Red, blue, and green represent haplotypes of Y900, Y2, and Y1, respectively; **f** Comparison of the nine different genotypes of *Ghd8* and *OsSPL13* in terms of HD and GYPP. Red represents haplotypes of Y900, and blue represents haplotypes of Y2 and Y1. Different letters above the bars represent significant differences between the groups (LSD) using one-way ANOVA (*P* < 0.05).

Although *LAX1* plays an important role in two-line hybrid rice, the better yield performance of the XI-GJ subspecies combination was likely due to the introduction of some super haplotypes, including *NAL1*, which is more common in GJ and is rarely used in XI (Huang et al., 2015; Xiao et al., 1995). We then simulated the combination of adding *NAL1* to *LAX1* and found that the GNPP and GYPP of the corresponding phase 3 super hybrid rice genotype combinations exhibited an increase, i.e., Y1 carried the heterozygous genotype of *LAX1* and the homozygous maternal genotype of *NAL1*, Y2 carried the heterozygous genotype of *NAL1* and the homozygous maternal genotype of *LAX1*, and Y900 carried the heterozygous genotypes of *LAX1* and *NAL1* (Figure 7e). This may partially explain the source of the difference in GNPP in the representative combinations of the phase 3 super rice. We also found among all nine genotype combinations, i.e., *LAX1* (homozygous maternal genotype) + *NAL1* (homozygous maternal genotype), the corresponding GNPP and GYPP reached 2130.4 and 33.4 g, respectively, which were still less than those of the double heterozygous genotype of Y900.

The rice blast resistance gene *RGA4* participates in the formation of resistance heterosis in young panicle and leaf tissues through different modes (DO and PDO) and is mainly regulated in *cis*, and its expression in leaves of R900 is relatively high. The allele bias in the hybrid gradually shifted from the young panicles in Y58S at the beginning to stems and leaves in R900 (Data S11), and QTN showed that R900 carried the haplotype that can increase the resistance to rice blast (Data S2). Further analysis showed that the protein homology between R900 and the *RGA4* donor was higher, and a similar phenomenon was also reflected in another resistance gene, *RGA5*, except that the overall expression level of *RGA5* was higher than that of *RGA4. RGA4* and *RGA5* are two adjacent *NBS-LRR* genes that are essential for leaf blast resistance of *Pia* (Okuyama et al., 2011) which may provide molecular evidence for the selective expression of favorable alleles in the growth and development of hybrids to enhance their disease resistance. This also shows that the heterosis formation for different traits in rice is dynamic, with temporal and tissue specificity.

## Discussion

Heading date is closely related to the high-yield and ecological adaptability, which is the key factor of rice heterosis. Although many loci of flowering time of rice had been cloned, their relationship and mechanism of hybrid vigor are still uncleared. The remarkable feature of R900 is superlong HD, and the corresponding hybrid rice Y900 also has the longest HD among the phase 3 super rice. In addition to *HD1*, the parents also have QTN differences in *Ghd7, HD17, Ghd8, OsCOL4, DTH2*, and other typical flowering control sites (Data S2). Population effect analysis showed that only *Ghd7* and *Ghd8* show the most significant effect on the delay of HD (Data S12). We analyzed other heterosis loci identified in this study and found that the grain type gene *OsSPL13* exhibits a *Ghd7*-like effect on HD (Figure 7b and Data S12), which has not been reported in previous studies (Si et al., 2016). The recently cloned grain regulatory site *GW3p6* (Wang et al., 2019) was identified as the gene *OsMADS1* that regulates the flowering stage, and *GW3p6* is a heterosis-related gene, which may provide a good reference for our subsequent studies. The effects of *Ghd8* separately combined with *Ghd7* and *OsSPL13* were further analyzed (Figure 7f), and the results show that the genotypic effects of the two combinations are broadly consistent, but the combination of *Ghd8* and *OsSPL13* may be slightly better than the other combination under the double-heterozygous genotypic effect; if the three genes are combined, the expected effects on flowering and yield accrual do not occur, so we speculate that there may be effects such as epistatic interactions between *Ghd7* and *OsSPL13*. These genetic effects of alleles can not only better explain why R900 and Y900 have longer HDs, but also provide new site selection for the fine-tuning of the flowering date and yield of rice.

Recently, Chen et al. (Chen et al., 2022) comprehensively reviewed the research progress of rice biology and functional genomics, including the new progress and future trends in the utilization of rice heterosis. Among them, the potential of heterosis between XI and GJ subspecies needs to be further explored. In this study, we found that GJ introgression and gene expression/regulation patterns are close connected (Figure 2a), and it is believed that as more important functional genes are cloned and analyzed, the mechanism involved in the heterosis formation will become increasingly clear.

## Methods

### Plant materials

Rice materials included the two-line hybrid rice Y1, Y2, Y900 and the corresponding restorer lines R9311, YH2, R900, and their common female parent Y58S. The F2 population was obtained from self-crossing of F1, Y900-F2 (379), and Y2-F2 (259) respectively. All the materials were planted and investigated traits in Changsha, China (112.93° E, 28.23° N). The leaves of R900 and Y58S were used for long-read, short-read, and Hi-C sequencing. Tissues such as 5mm and 10mm panicle length, flag leaf at heading, and stem under the panicle were used for RNA sequencing, with 36 samples for each tissue (three individual plants and three biological replicates). The primers used to verify the expression levels of heterosis-related genes were listed in Table S13.

### Genome assembly and annotation

For NGS short reads, library construction and sequencing were performed on the MGISEQ-2000 platform (BGI, Shenzhen, China). For long reads, this was done on the PacBio Sequel II platform. Genome assembly was performed using Canu v1.8 (Koren et al., 2017), followed by polish using Pilon (Walker et al., 2014). The 144.7 million and 148.8 million paired-end clean reads generated by Hi-C sequencing in R900 and Y58S, respectively (Table S3), were clustered, sorted, and positioned in contigs using Lachesis. Based on the embryophyta_odb10 database, BUSCO (Simao et al., 2015) was used to evaluate the genome integrity. LAI was used as the standard for assessing the assembly of repetitive sequences (Ou et al., 2018). The NGS reads were aligned to the assembled genome using the BWA-MEM algorithm (Li and Durbin, 2009). The collinearity alignment between rice genomes was performed using Minimap2 (-x asm5) (Li, 2018).

Repeat sequences were masked using RepeatMasker (Zhi et al., 2006) and TE library from Repbase library (Edition-20181026) (Bao et al., 2015). Genes were predicted by integrating evidence from ab initio, homology-based, and mRNA. Specifically, Augustus (Stanke et al., 2006) and GlimmerHMM (Majoros et al., 2004) were used to perform de *novo* gene prediction on genomes that had been masked for repetitive sequences; homology alignment by exonerate for all known plant proteins in the UniProt database and predicted proteins in rice (MSU7), maize (B73 v5) and Arabidopsis (TAIR10); and BLAT (Kent, 2002) and GMAP (Wu and Watanabe, 2005) for all *Oryza sativa* mRNAs from the GenBank database and transcripts assembled by Trinity (Grabherr et al., 2011) for four tissues in this study. The above-predicted gene models were integrated using EVidenceModeler (Haas et al., 2008) and finally updated using PASA (Haas et al., 2003). TRNA, rRNA, snRNA, and miRNA were predicted by using tRNAscan-SE (Schattner et al., 2005), Barrnap, and INFERNAL (Nawrocki and Eddy, 2013). Functional annotation of protein sequences using BLASTP alignment with filtering condition *E* value < 1e^-5^. TE-related genes were annotated in the same way as Zhang et al. (Zhang et al., 2016). Fisher’s exact test was used to detect the GO terms and KEGG pathways with significant gene enrichment.

### Genomic variant identification and validation

Y58S was used as the reference genome, and MUMmer v4.0 (-maxmatch -c 90 -l 40) (Marcais et al., 2018) was used to align with the R900, R9311, R498, NIP, HZ, TF, MH63RS1, and ZS97RS1 genomes, respectively. delta-filter -m filtered the results, show-snps (-Clr th) identified SNPs and Indel. SnpEff (Cingolani et al., 2012) annotated the variants. The density distribution of SNPs and Indel on the genome was calculated with a sliding window of 100 kb size. Based on the MUMmer results, the default parameters of SyRI Pipeline (Goel et al., 2019) were used to identify structural variants between genomes. The variants detected by SyRI include two types: genomic rearrangements and sequence variants. These variants were converted into three SV types, PAV, inversions and translocations, according to the definition of SyRI results. CPL, DEL, DUP/INVDP (loss), HDR, NOTAL and TDM in the Y58S genome were defined as Absence SVs (relative to Y58S). CPG, INS, DUP/INVDP (gain), HDR, NOTAL and TDM in the query genome were defined as Presence SVs (relative to Y58S). INV is considered inverted SVs. Both TRANS and INVTR are considered translocated SVs.

For large structural variants, NGS was aligned to the Y58S and NIP genomes using BWA MEM (Li and Durbin, 2009), and the sequenced bam files were imported into IGV (Integrative Genomics Viewer) (Robinson et al., 2011) and verified by the variant locus reads coverage to confirm the genotypes of different varieties. Rice varieties included Y58S, R900, R9311, R498, NIP, HZ, TF, MH63RS1, ZS97RS1, and YH2, with NGS data from previous studies (Lv et al., 2020). The XI-GJ genetic composition and introgression bin maps were constructed using the method described by Chen et al. with 3K-RG as the background and a 100-kb size sliding window between Y58S, R900, R9311, YH2 and NIP using high-density SNPs (Chen et al., 2020).

### RNA-sequencing data analysis

Using Y58S as the reference genome, the “new Tuxedo” protocol (Pertea et al., 2016) was used for the analysis of assembly of transcripts, quantification of gene expression levels, etc., and DESeq2 (Love et al., 2014) for differentially expressed gene analysis based on grouping information. Differentially expressed genes were defined as Fold Change > 2 and FDR < 0.05. Transcript per million (TPM) was calculated to represent gene expression. 0 < TPM <= 1 was defined as low expression. For specifically expressed genes, more than 2/3 of the samples must have TPM > 0 to be considered as expressed.

Based on the differential gene expression trends of R900, Y58S, and Y900 in individual tissues, the heterosis patterns were classified according to three different modes (Mode I-III) (Figure 4b). Mode I assigned all possible gene expression patterns to 12 (Swanson-Wagner et al., 2006). Mode II based on 12 patterns of F1 and parental gene expression can be summarized into five patterns, namely H2P, CHP, B2P, CLP, and L2P represent higher than both parents, close to higher parent, between both parents, close to lower parent, and lower than both parents (Wei et al., 2009). Mode III further classified the five models into partial dominant (PDO), dominant (DO) and over dominant (ODO) according to the definition of heterosis.

### ASE analysis

First, the transcriptomic libraries of all samples were aligned to the R900 and Y58S genomes using HISAT2 (Kim et al., 2019), then headers were added to the bam files using Picard, and SNPs were identified using the GATK pipeline (McKenna et al., 2010). i.e. MarkDuplicates to remove duplicates, SplitNCigarReads to remove reads in the intron region, HaplotypeCaller to detect variants, and VariantFiltration to filter variants. Further, the ASEReadCounter program (-min-depth 8) was used to identify the allele origin of each sample separately and filter out the SNPs that did not meet the requirements (the number of reads from samples of the same species as the reference genome should be much smaller than the number of reads from samples of different species). To eliminate the preference for a single genome, we combined all SNPs between the two genomes and the SNPs identified to R900 and Y58S as the reference genome, respectively. A total of six replicates of the three genotypes per tissue were used to finally identify ASEGs by comparing the difference in the number of reads of the two alleles in F1 by DESeq2 (*P* adjust < 0.05) (Love et al., 2014).

### Analysis of *cis-* and *trans*-regulated genes

Referring to the methods of previous studies (Bao et al., 2019; Wittkopp et al., 2004), *cis-* and *trans-*regulated genes were identified by comparing the gene expression of the two parents and the two alleles in F1 (Figure 6a).

### Data access

All the raw data generated for this project are archived at the NCBI BioProject database under accession number PRJNA825106 (*<u>Reviewer’s link:</u> https://dataview.ncbi.nlm.nih.gov/object/PRJNA825106?reviewer=29koia385tcvp5qsg34mgp6j5v*) and the National Genomics Data Center BioProject database under accession number PRJCA009056. The raw sequencing data for the genome and transcriptome are deposited in the Genome Sequence Archive database under accession numbers CRA006614 (*<u>Reviewer’s link:</u> https://ngdc.cncb.ac.cn/gsa/s/l22q9A04*) and CRA006615 (*<u>Reviewer’s link:</u> https://ngdc.cncb.ac.cn/gsa/s/6K9948j9*). The R900 and Y58S genome assembly sequences are deposited in the Genome WareHouse database under accession numbers GWHBJTR00000000 (*<u>Reviewer’s link:</u> https://ngdc.cncb.ac.cn/gwh/Assembly/reviewer/POvTmMCdZCxbhErJeWJLTfohbbadxOwrPzPUZZKXEfFfuDxGyLEGmLWbMKLLAUoO*) and GWHBJTS00000000 (*<u>Reviewer’s link:</u> https://ngdc.cncb.ac.cn/gwh/Assembly/reviewer/MFQSNFjhNAokZQNXkJuXNppWXqdHABfDIOEjmmylTubIMwlwbjuMurDzXeCUKFRq*). The supplemental data can be accessed from the FigShare database with DOI: 10.6084/m9.figshare.20140235 (*<u>Reviewer’s link:</u> https://figshare.com/s/a7e0e1fb46480aca0835*).

## Acknowledgements

This work was supported by the National Key Research and Development Project (2016YFD0101100), the Science and Technology Innovation Program of Hunan Province (2021NK1001, 2021RC4066, 2021NK1003, 2021NK1012, and 2021RC3113), Major Science and Technology Program in Hainan Province (ZDKJ2021002), National Natural Science Foundation of China (U21A20208 and 31801341), Natural Science Foundation of Hunan Province (2020JJ4456). We thank Qiusheng Xu (State Key Laboratory of Hybrid Rice, Hunan Hybrid Rice Research Center, Hunan Academy of Agricultural Sciences) for their useful discussion and opinion on the article.

## Author contributions

D. Yuan., and B. Wang. conceived the research project and designed the experiments. Z.S. managed the project. Z.S. and J. Peng wrote the manuscript. D. Yuan., B. Wang., Z.S., J. Peng., and Q.L. participated in the article discussion and performed the bioinformatics work. Z.S., J.D., S.C., Q.H., J.W., Y.T., M.D., D.Y., Y.T., X.S., J.C., X.S., L.L., R.P., H.L., T.Z., and N.X. conducted the experiments.

